# Optimizing dsRNA sequences for RNAi in pest control and research with the dsRIP Web-Platform

**DOI:** 10.1101/2025.01.09.631884

**Authors:** Doga Cedden, Gözde Güney, Michael Rostás, Gregor Bucher

## Abstract

**Background:** RNA interference (RNAi) is a tool for studying gene function and has emerged as a promising eco-friendly alternative to chemical pesticides. RNAi relies on delivering double-stranded RNA (dsRNA), which is processed into small interfering RNA (siRNA) to silence genes. However, so far, knowledge and tools for optimizing the dsRNA sequences for maximum efficacy are based on human data, which might not be optimal for insects and pest control.

**Results:** Here, we systematically tested different siRNA sequences in the red flour beetle *Tribolium castaneum* to identify sequence features that correlated with high efficacy using pest control as a study case. Thermodynamic asymmetry, the absence of secondary structures, and Adenine at the 10^th^ position in antisense siRNA were most predictive of insecticidal efficacy. Interestingly, we also found that in contrast to results from human data, high, rather than low GC content from the 9^th^ to 14^th^ nucleotides of antisense was associated with high efficacy. Consideration of these features for the design of insecticidal dsRNAs targeting essential genes in three insect species improved the efficacy of the treatment. The improvement was associated with a higher ratio of the antisense, rather than sense, siRNA strand bound to the RNA-induced silencing complex. Finally, we developed a web-platform named dsRIP (https://dsrip.uni-goettingen.de), which offers tools for optimizing dsRNA sequences, identifying effective RNAi target genes for pest control, and minimizing risk to non-target species.

**Conclusions:** The identified sequence features and the dsRIP web-platform allow optimizing dsRNA sequences for both application of RNAi for pest control and research.

## Introduction

Crop production must increase to feed the growing human population that is estimated to reach ∼10 billion people in the upcoming three decades [1, 2]. Pests and diseases account for 20-40% of losses in major crop yields [3], and several insect species act as vectors for serious human diseases [4]. Currently, the management of crop pests and human-disease vectors is heavily dependent on the application of chemical pesticides, such as neonicotinoids [5]. Sole reliance on pesticides with limited mode of action repertoire has led to the emergence of pesticide-resistant populations, leading to ineffective control of target pest and vector populations [6–8]. Moreover, chemical pesticides are also a threat to non-target organisms such as honeybees and natural enemies (e.g., predators and parasitoids of pests) due to their non-species-specific characteristics [9–11]. Hence, there is both biological and societal pressure to adopt more sustainable and eco-friendly management strategies.

Recently, RNA interference (RNAi) has emerged as a promising eco-friendly alternative to chemical pesticide applications [12–15]. The RNAi pathway is triggered by double-stranded RNA (dsRNA), which can enter insect cells through endocytosis and is released into the cytoplasm via the retrograde and endoplasmic-reticulum-associated protein degradation pathways [16, 17]. Cytoplasmic dsRNA is processed by the RNase III enzyme Dicer-2 into ∼21 nucleotide-long small interfering RNA (siRNA) duplex with characteristic 2-nucleotide (nt) overhangs at the 3’ ends [13, 18]. The siRNA duplex is loaded onto an Argonaute-2 enzyme, forming the RNA-induced silencing complex (RISC) [19]. The RISC cleaves one of the strands of the siRNA duplex (the passenger strand) and retains the other (the guide strand). Subsequently, RISC finds and cleaves mRNA targets that are complementary to the guide strand, thereby leading to sequence-specific gene silencing [20].

In the context of pest management, genes with essential functions are targeted through the cultivation of dsRNA-expressing crops (e.g., SmartStax PRO maize with dsSnf7 against Western corn rootworm *Diabrotica virgifera virgifera*) [21] or sprayable dsRNA formulations (e.g., Ledprona with dsPSMB5 as active ingredient against Colorado potato beetle *Leptinotarsa decemlineata*) [22]. Hence, the first step in the establishment of an RNAi-based strategy against a pest is the identification of highly effective target genes, i.e., those causing the highest and quickest mortality following the delivery of minimal dsRNA doses [23, 24]. Recently, the genome-wide RNAi screen in *Tribolium castaneum* established a list of highly effective target genes, and a subset of these genes were also highly effective targets in multiple leaf beetles (Chrysomelidae) [25, 26]. Hence, the problem of effective target gene selection has been, for the most part, solved, especially for coleopteran pests. However, the parameters for designing a dsRNA optimized for maximum efficacy beyond the selection of the target gene remain unclear.

A potential frontier in enhancing RNAi-based pest control is the rational designing of the dsRNA sequence to effectively silence the target genes. A typical mRNA is around 2000 bp in length [27], while pesticidal dsRNA delivered either via transgenic crops or foliar application are between 200-500 bp [22, 28], leaving a notable degree of freedom for the design of the dsRNA. The only well-established finding that is widely used during dsRNA designs for pest control concerns the length, which should be at least 60 bp for efficient cellular dsRNA uptake [28–30]. Nonetheless, there are indications from the literature that adopting a more nuanced approach during the designing of the dsRNA sequence might increase the pest control efficacy by improving the RNAi response. Although long dsRNA is used in pest control, the main mediators of the RNAi response are the siRNA processed from the dsRNA. Notably, efforts in developing therapeutic siRNA against human diseases heavily focused on the identification of siRNA features for optimized gene knock-down [31, 32]. The findings converged on selected features, such as the thermodynamic asymmetry of the siRNA duplex, that are used to guide the selection of effective siRNA candidates for silencing gene of interests in human cells [33, 34]. Based on the siRNA efficacy parameters identified in human cells, several software, including DEQOR and siDirect, were developed to find sequence stretches that contain effective siRNAs against target genes [33, 34].

Further work also demonstrated the conserved mechanistic basis for the importance of these siRNA features. For instance, the strand with the weakly paired 5’ end in the siRNA duplex is preferentially selected by the RISC as the guide strand due to the thermodynamic asymmetry sensing capability of dsRNA binding helper proteins (e.g., PACT in humans and R2D2 in *Drosophila melanogaster*), enabling the possibility to bias the guide RNA selection in favor of the antisense strand rather than the sense strand [35–38]. Also, siRNAs with weaker 5’ end were higher in abundance after the delivery of an insecticidal dsRNA to the *Psylliodes chrysocephala*, cabbage stem flea beetle [39]. Furthermore, higher insecticidal efficacy of dsRNA region containing more siRNA predicted by siRNA design algorithms [40] suggested that insecticidal dsRNA designs could be enhanced by considering siRNA sequence features. Moreover, the fact that insects lack the secondary siRNA generation mechanism [41, 42] described in *Caenorhabditis elegans* [43], suggests that insects rely on the siRNA pool processed directly from the uptaken dsRNA. This implies that the sequence of the delivered dsRNA is of great importance in mounting an effective RNAi response and achieving high insecticidal efficacy in insect pests. Algorithms based on human data, such as DEQOR [34], were used to select the putatively best regions within a target sequence for insects [25]. However, how far these predictors based on human data can be transferred to improve the insecticidal efficacy of dsRNA has never been systematically tested in insect pests. Also, currently there are no publicly available tools that guide designing dsRNA based on siRNA features against effective target genes while minimizing off-target effects on non-target organisms.

In this study, we first wanted to identify the parameters relevant to insect siRNAs using beetles as model. Therefore, we systematically tested the insecticidal efficacy of individual siRNA sequences embedded in a non-targeting dsRNA backbone targeting the same essential gene in *T. castaneum*. We found that similar to human data, features including thermodynamic asymmetry and lack of secondary structures were predictive of high efficacy. Some factors known from humans were not important in our tests, including mRNA accessibility. Interestingly, higher GC content between 9^th^ and 14^th^ nucleotides was associated with high efficacy, rather than lower GC content as found in human cells. Based on our findings, we optimized the dsRNA design against multiple essential gene targets in *T. castaneum* and in two leaf beetles. Moreover, we sequenced RISC-bound small RNA following the delivery of dsRNA to further understand the reason behind the differences in the insecticidal efficacies of dsRNA. We found that in optimized dsRNAs, a higher ratio of the antisense siRNA was bound to RNA-induced silencing complex. Lastly, we integrated our parameters into an online dsRNA optimization algorithm that helps designing effective dsRNA for use in pest control and basic science independent of the species. Further, we added tools that guide the selection of effective RNAi target genes in diverse pests and minimize detrimental risk to non-target organisms. This open-source tool was made available as a web-based platform named “Designer for RNA Interference-based Pest Management”, shortly dsRIP.

## Results

### Identification of insecticidal efficacy associated siRNA features

First, we wanted to empirically obtain a set of siRNA sequences that act with different efficacies in RNAi. To that end, we tested the insecticidal efficacy of randomly selected 31 siRNAs (21-nt long), each targeting different regions of the same essential gene (*Tc-gawky*; see Supporting File 1: *T.castaneum* siRNA insert for sequences). The siRNAs were inserted individually into a non-targeting backbone (mGFP), leading to a dsRNA template of 232 bp length (Fig. 1a). First, we wanted to confirm that the respective dsRNA was processed in injected larvae and that both the siRNA to be tested and those from the backbone were detectable as RISC-bound siRNA. Indeed, RISC-bound small RNA-seq showed a non-random processing along the entire dsRNA *in vivo* (see Fig. 1a). We noted that the reads corresponding to the tested siRNA were not the dominant peak among all generated siRNA. Interestingly, the profile of RISC-bound siRNAs from the backbone was highly reproducible in two different experiments (compare the two plots in Fig. 1a, n = 1). Hence, the artificial dsRNAs are indeed processed, and the generation of RISC-bound siRNAs is a non-random process.

**Figure 1.**
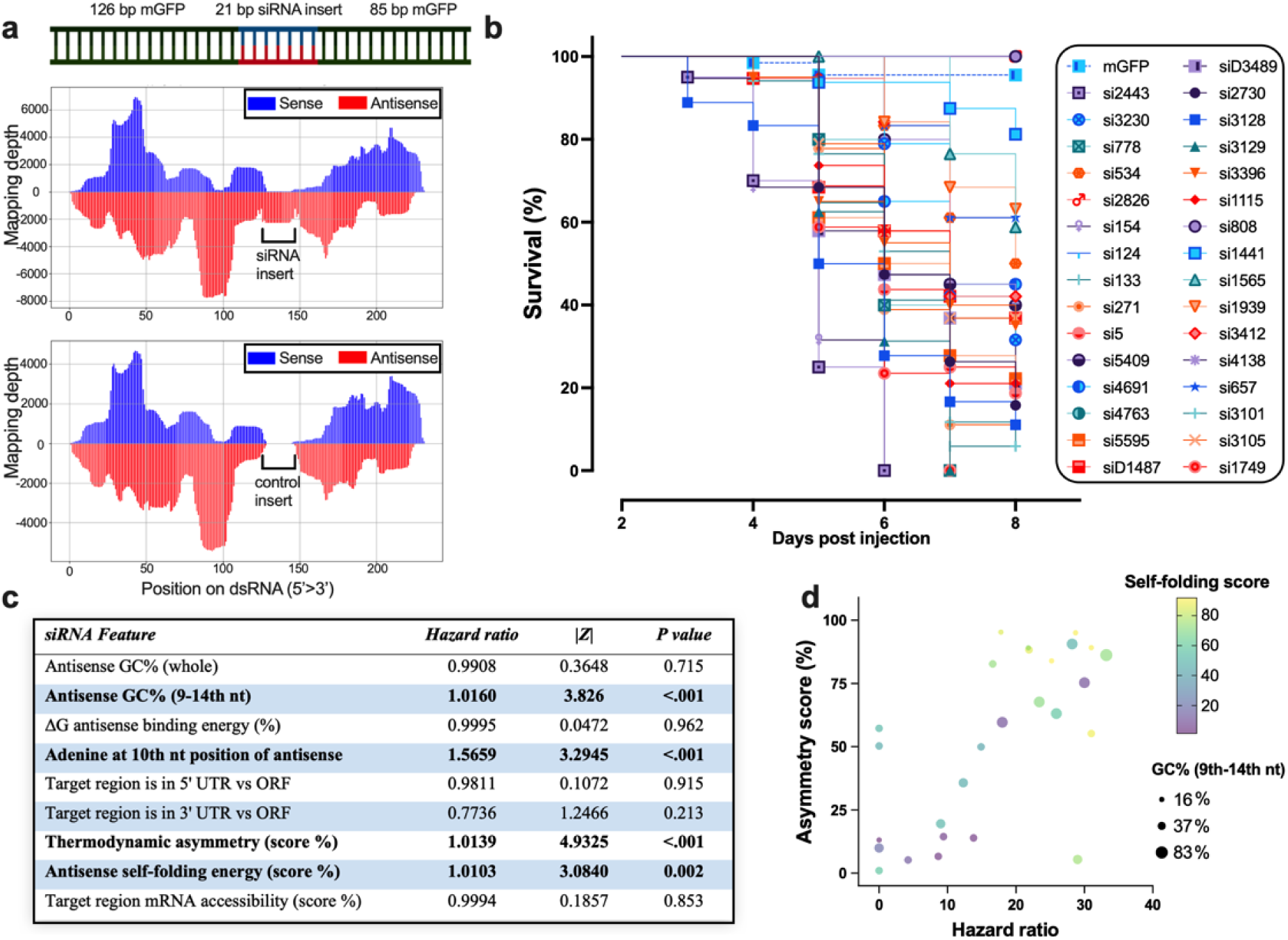
Testing the efficacy of different siRNAs in *Tribolium castaneum*. (a) Validation of the processing of siRNA inserted dsRNA in *T. castaneum*. An siRNA of 21 nt targeting an essential gene was inserted into a backbone of mGFP sequences (GFP sequence optimized for no off-targets). Prior to the bioassays, we confirmed that the target siRNA was processed into the RISC complex along with the non-targeting siRNAs after injection of 200 ng/ul per L5 larvae (n = 1 representing 5 injected larvae). The upper plot shows the mapping depth of 21-nt RISC-bound siRNA perfectly mapping onto the *Tc-cdk9* (TC003392) targeting siRNA and backbone dsRNA. The plot below shows the mapping on to the same dsRNA template when larvae were injected with a dsRNA with another siRNA insert that was targeting *Tc*-*gyc32e* (TC011636). Note the very similar profile of backbone siRNA processing in these two independent experiments. (b) L5 larvae (n = 20 per group) were injected with 1000 ng of dsRNA containing one siRNA, which was complementary to the different regions of the essential gene Tc-*gawky, respectively* (see supporting file 1 for details). The Kaplan-Meier survival curves were plotted for each siRNA along with the non-siRNA inserted dsmGFP as a control. (c) To reveal the sequence features that predict efficacy, the survival data for each siRNA was fit with respective siRNA sequence features into a Cox proportional-hazards model to determine the main effects of siRNA sequence features on the relative hazard. The scores that significantly predict efficacy are shown in bold. (d) The hazard ratio for each siRNA insert group was calculated according to the log-rank method and plotted against three continuous variables that had a significant effect on insecticidal efficacy according to the Cox regression analysis.

To quantify the insecticidal efficacies of these 31 siRNAs in bioassays, we injected dsRNA with a concentration of 1 µg/µl into L5 *T. castaneum* larvae (n = 20). The bioassays showed that dsRNAs with different siRNAs had different impacts on the larval survival, ranging from 100% lethality after 6 days to 20 % lethality after 8 days (Fig. 1b). To find predictors of efficacy, we determined values for nine sequence-based features of each siRNA insert (see Fig. 1c), which had been identified as being important in previous work based on human data^49^. These features include the thermodynamic asymmetry score, where a high score indicates that the 5’ end of the antisense strand forms weaker bonds; the antisense self-folding energy score, where a high score reflects the absence of strong self-folding structures; and the mRNA accessibility score, where a high score signifies that the siRNA targets a more accessible region in the mRNA^49^. Next, we used the Cox regression method to identify predictors of siRNA insecticidal efficacy. The Cox regression model (Harrell’s C-statistic = 0.76 with 95% confidence interval of 0.73 to 0.79) indicated that four of the nine features were significantly associated with increase in insecticidal efficacy (P < 0.05, Fig. 1c-d): Increase in thermodynamic asymmetry score, antisense self-folding energy score, % GC content from 9^th^ to 14^th^ nt of the antisense strand and the presence of an Adenine at the 10^th^ nt position of the antisense strand. In contrast to previously established features identified in human cells^49^, the target mRNA region (i.e., ORF vs UTR); the mRNA accessibility score, % GC of the entire antisense strand, and binding energy of the antisense strand to the target mRNA were not significant predictors of efficacy (P > 0.05, Fig. 1c). Overall, the bioassays with siRNA inserted dsRNAs suggested that specific sequence features of siRNAs can affect the insecticidal efficacy and therefore need to be taken into consideration when designing dsRNA fragments. Further, our results indicate that only some parameters known from human data are transferrable to insects. This shows that an insect-specific assessment is advisable when aiming to optimize dsRNA efficacy.

### Testing siRNA feature-based dsRNA designs

The observation that siRNAs exhibited different efficacies suggested that long dsRNA could be optimized for higher insecticidal efficacy by enriching for siRNAs with these features. To that end, we selected 8 genes previously identified as very effective target genes through genome-wide RNAi screen in *T. castaneum* [25] and designed 2 dsRNA, respectively, with either high or low mean siRNA efficacy score. We used the significant predictors of efficacious siRNAs and their coefficients (see Fig. 1c and [Z] scores therein) to calculate an siRNA efficacy score for all possible siRNAs that can be processed from the target mRNAs. Next, we implemented a sliding window that identified the 300 bp region within the target gene sequence that had the highest mean siRNA efficacy score within the ORF. The sliding window calculates the mean of siRNA efficacy scores for every possible siRNA that could be generated from the window and then shifts the window one nucleotide at a time, eventually finding the window with the highest mean score. Using the same calculation, a 300 bp region with the lowest mean siRNA score within ORF was selected for comparison. In addition, we also tested the region with the highest mean siRNA score without restricting the prediction to the ORF. Finally, we tested the region with the highest mRNA accessibility score without restricting the prediction to the ORF (the same calculation as in Fig. 1 was used, and the mean was calculated for every possible window). dsRNAs not restricted to ORF were not tested if they majorly overlapped (>60%) with any of the other tested regions.

In the tests with most target genes, namely, *Tc-gawky* (Fig. 2a), *Tc-hr3* (Fig. 2b), *Tc-klp61F* (Fig. 2c), *Tc-nito* (Fig. 2d), *Tc-eIF3a* (Fig. 2e), and *Tc-rpt7* (Fig. 2g) the dsRNA fragments with high mean siRNA efficacy score performed better than dsRNA with low mean siRNA efficacy score (both within ORF). For statistical testing of this observation, we used a multilinear regression model to analyse the contributions of different score types to the dsRNA insecticidal efficacy (Tab. S1). To that end, we tested the effects of siRNA efficacy score, mRNA accessibility score, the target region in mRNA (ORF or UTR), and target genes on the insecticidal efficacy (i.e., hazard ratio of dsRNA treatment group over dsmGFP control group) upon dsRNA injection. The model had a multiple R-value of 0.79 (df = 17) and passed the Shapiro-Wilk normality test (P = 0.84). The bubble plot reveals that a main predictor remains the selection of the target gene (see Tc-*gawky* and Tc-*nito* with the highest hazard ratio). Interestingly, including the UTRs for searching allows finding fragments with a higher siRNA efficacy score (purple dots in the upper part) but this led to no increase in hazard ratio. Overall, the siRNA efficacy score predictions correlated very well with the measured insecticidal efficacy (Fig. 2j). The results of the statistical test confirmed that the siRNA score of the dsRNA was the most significant and positive predictor of insecticidal efficacy (1.462 increase in hazard ratio per 1 increase in siRNA score, P = 0.003, Fig. 2j). mRNA accessibility score was not a significant predictor of insecticidal efficacy (P = 0.41, Fig. 2j), which is in line with our observation in the bioassays testing single siRNAs (Fig. 1c). In contrast to the single siRNA data, however, targeting of the ORF was associated with higher insecticidal efficacy compared to the targeting of UTR (9.91 increase in hazard ratio, P = 0.011, Fig. 2j). This may be because the combined effect of multiple siRNAs targeting the ORF as part of a long dsRNA fragment creates a statistically significant difference, whereas the impact of a single siRNA remains below the significance threshold. As expected, targeting different target genes had a major impact on the insecticidal efficacy. For instance, targeting of *Tc-rpt1* was less effective compared to the targeting of *Tc-gawky* (15.83 decrease in hazard ratio for *Tc-eIF3a* compared to the reference *Tc-gawky*, P = 0.005, Fig. 2j). Overall, the analysis shows that choosing the best target gene and the dsRNA region with the best siRNA efficacy score within the ORF of that target gene improved insecticidal efficacy in most cases. They also show that the dsRIP algorithm is a very good predictor and can benefit RNAi-based pest control strategies. Hence, we implemented the algorithm used for that purpose in dsRIP, a new online platform to predict efficient dsRNA sequences with high insecticidal efficacy.

**Figure 2.**
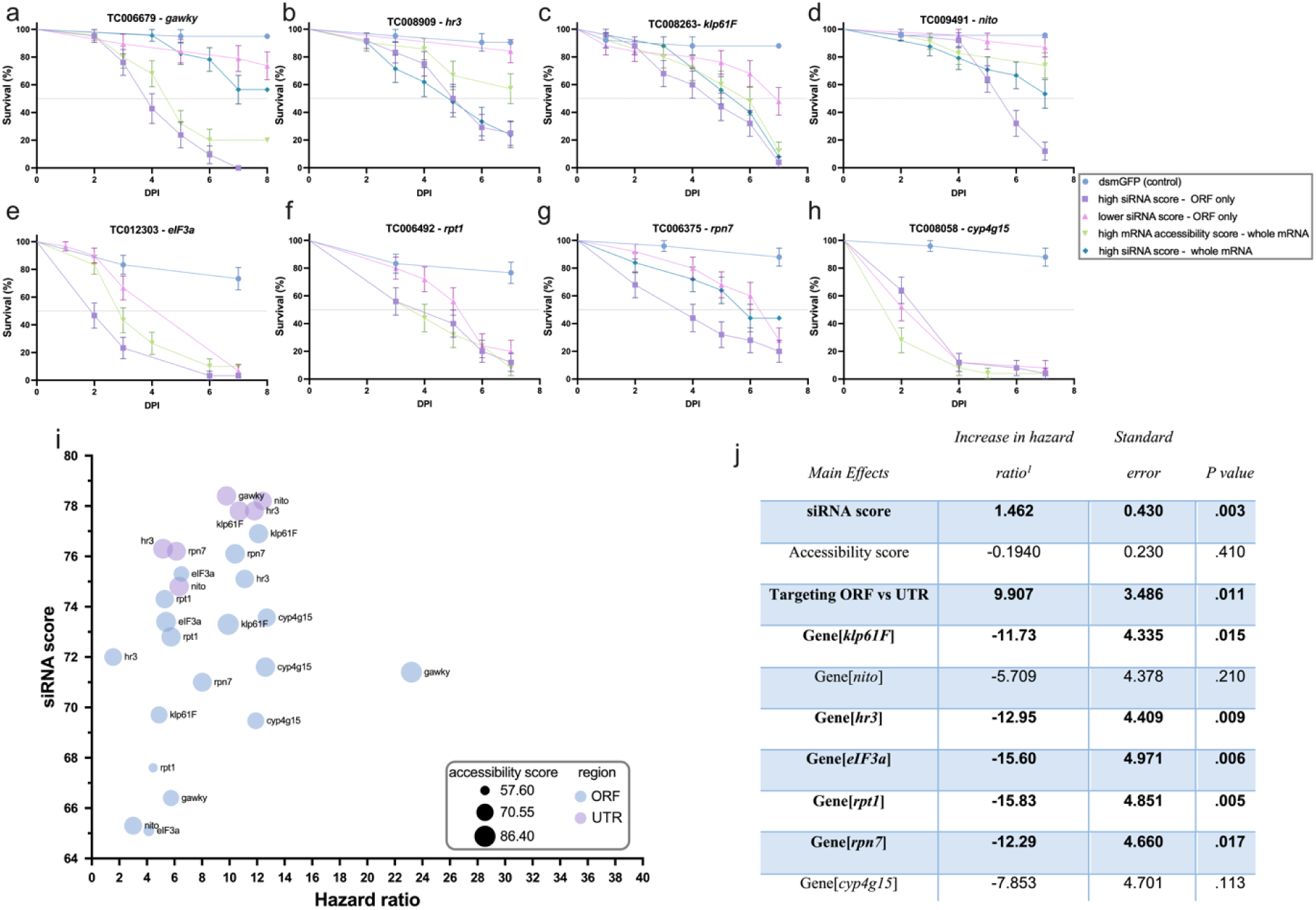
Testing the efficacy of dsRNAs designed using dsRIP in *Tribolium castaneum.* Several different dsRNA predictions were tested in *T. castaneum* via the injection of 1 ng/µl dsRNA targeting one essential gene, respectively, into L5 larvae (n = 25). Three or four different dsRNA were tested for insecticidal efficacy against 8 genes, namely, Tc-*gawky* (a), *Tc-hr3* (b), *Tc*-*klp61F* (c), *Tc*-*nito* (d), *Tc*-*eIF3a* (e), *Tc*-*rpt1* (f), *Tc*-*rpt7* (g), *Tc*-*cyp4g15* (h) in parallel to control dsmGFP injected larvae. The survival curves (mean survival ± standard error) showed that dsRIP optimization for siRNA efficacy score indeed increased the lethality of the treatment. (i) Using survival data for all tested dsRNAs, the hazard ratio was calculated against the control dsmGFP group. The bubble plot shows that the selection of target genes remains key (compare *Tc-gawky* and *Tc-nito* to the right). It also indicates that the higher siRNA efficacy scores that were obtained by including UTRs did not result in increased lethality (see purple dots in the top part). (j) All data was fitted into a multiple regression model to investigate the main effects on the hazard ratio of the mean siRNA efficacy and mRNA accessibility scores, of restricting the target region to ORF and of choosing the target gene. The model suggested that the siRNA efficacy score could be used to determine the dsRNA target region with the highest insecticidal efficacy. Also, targeting the ORF and selecting the target gene turned out to be the key parameters associated with insecticidal efficacy. P values < 0.05 were accepted as significant contributors to the hazard ratio and indicated in bold. *Tc*-*gawky* was taken as the reference level for the target gene factor, compared to which *klp61F, hr3, elF3a, rpt1* and *rpn7* were significantly worse choices.

### High scoring dsRNA have higher RISC-bound antisense siRNA strand

To understand the reason behind the insecticidal efficacy improvement of dsRNA with high siRNA score, we performed RISC-bound sRNA-seq in dsRNA-injected *T. castaneum* larvae. As thermodynamic asymmetry was a major siRNA feature used during the selection of the optimized dsRNA region, we hypothesized that delivery of optimized dsRNA might be leading to a higher RISC-bound antisense/sense strand ratio.

The dominant siRNA length was 21 nt (Fig. S1) [39, 44], the same as the siRNA length used in the individual siRNA testing bioassays (Fig. 1). We observed a diverse pool of both sense and antisense siRNA mapping onto the respective dsRNA in all treatments (see blue (sense) and red (antisense) bars in Fig. 3a-h). Of note is that only antisense siRNA strands can lead to target mRNA destruction. Confirming our hypothesis, the antisense/sense siRNA strand ratio was in most cases (7 out of 8 gene targets) higher with dsRNA with higher siRNA efficacy scores (P < 0.018 by paired t-test, Fig. 3i). On average, they produced 20.1% more antisense siRNA compared to the lower scoring dsRNA targeting the same essential gene (Fig. 3i). Interestingly, in *Tc-cyp4g15* the higher scoring dsRNA did not improve the antisense/sense siRNA ratio (Fig. 3f), but that correlated with the bioassays, where the insecticidal efficacy was not improved (Fig. 2h).

**Figure 3.**
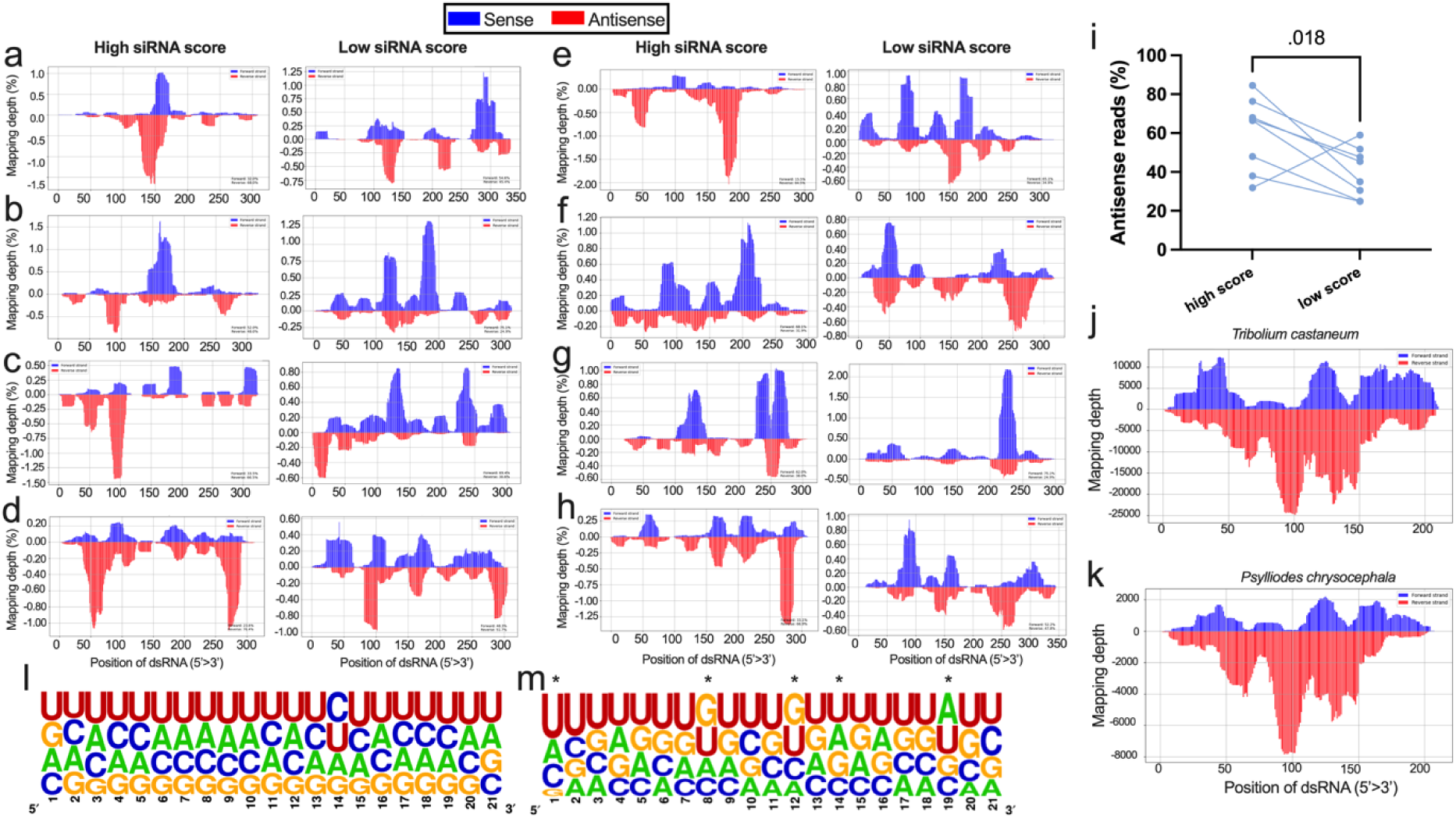
RISC-bound sRNA-seq following dsRNA delivery. (a-h) dsRNA with high and low siRNA efficacy scores against essential genes were injected, and subsequently, RISC-bound siRNAs were sequenced. Targeting siRNAs (i.e. antisense) are shown in red, while non-targeting (i.e. sense) siRNAs are shown in blue for eight different genes: Tc-*gawky* (a), Tc-*hr3* (b), Tc-*klp61F* (c), Tc-*rpn7* (d), Tc-*eIF3a* (e), Tc-*cyp4g15* (f), Tc-*nito* (g) and Tc-*rpt1* (h) (5 larvae pooled into n = 1, see Fig. S2 for reproducibility of sRNA-seq experiments). The processing is not uniform and the highly efficient dsRNA fragments seem to generate more RISC-bound antisense siRNAs (see increased red bars) (i) Statistical test confirmed that in dsRNAs with high siRNA efficacy score, the percentage of antisense siRNA strand reads is increased (paired t-test-connecting line indicates the same target gene). (j,k) siRNA processing is very similar in different beetles. The mapping of the control dsRNA (dsmGFP) after its delivery to *T. castaneum* (larval injection; panel j) was compared to *Psylliodes chrysocephala* (oral feeding of dsRNA, panel k). This suggests that our findings in *T. castaneum* can be transferred to other insect species. (l,m) The nucleotide frequencies in the least abundant (bottom ten percentile, panel l) and most abundant (top ten percentile, panel m) siRNAs were compared, with asterisks indicating notable differences between the two. Abundant siRNAs tend to be U-rich and G-poor at the 5’end and more G-rich in the middle of the strand, especially in the 8^th^ and 12^th^ positions.

In order to find sequence motifs enriched in RISC-bound siRNA strands, we compared the sequences of the antisense siRNAs from all tested genes with low abundance (bottom 10%, Fig. 3l) with those with high abundance (top 10%, Fig. 3m). There was a bias in favour of Uridine and against Guanine at the 5’ end of highly abundant 21 nt antisense siRNAs. Such a bias indicates thermodynamic asymmetry, where 5’ end of the siRNA forms weaker bonds. Further, we found that highly abundant siRNA contained Guanine more frequently in the middle of the strand, especially in the 8^th^ and 12^th^ positions (Fig. 3m).

To test the transferability of insights from *T. castaneum* to other insects, we performed RISC-bound sRNA-seq after the oral delivery of the control dsRNA, dsmGFP, to *Psylliodes chrysocephala* adults in parallel to larval injection into *T. castaneum* of the same dsRNA sequence (Fig. 3j-k). We observed a very similar RISC-bound siRNA abundance pattern in both insects, suggesting dsRNA processing and guide strand selection mechanisms are very similar, at least between these two beetles.

Overall, these RISC-bound sRNA-seq experiments suggested that the higher efficacy of dsRNAs may at least in part be due to the increased portion of antisense siRNAs that are incorporated into RISC.

### Application of the identified parameters to enhance RNAi-mediated pest control

To test the capability of the dsRNA designing strategy established in this study, which will be referred to as dsRIP, we selected *Psylliodes chrysocephala* and *Leptinotarsa decemlineata* to compare the performance of dsRNA designed by dsRIP with dsRNA used in previous studies. Three dsRNAs targeting the *P. chrysocephala* proteasome had been identified as highly insecticidal before [26]. We used dsRIP to design an improved fragment and tested both versions in parallel. Notably, the dsRNA designed by dsRIP showed significantly improved insecticidal efficacy compared to the dsRNA used in the previous study targeting the same gene (P < 0.05 by log-rank, hazard ratios: 1.8-2.5, Fig. 4a). Interestingly, the highest improvement in hazard ratio (2.5) was observed for the dsProsB7 which had the least insecticidal efficacy among the dsRNA tested in the previous study. These results show that choosing an optimized fragment is of key importance for high insecticidal efficacy. To test for sublethal effects, we also checked the feeding inhibitory effects of dsRNA (Fig. 4c). dsRNA designed by dsRIP significantly improved the feeding inhibitory effects measured 5 days post-treatment in the case of targeting Pc-*rpt4* or Pc-*prosB7*, but not Pc-*rpt1* (P < 0.05 by Šídák’s test). Note that the feeding behaviour peaks around 5 days post-adult emergence in control CSFB (Fig. 4c), as described previously [26].

**Figure 4.**
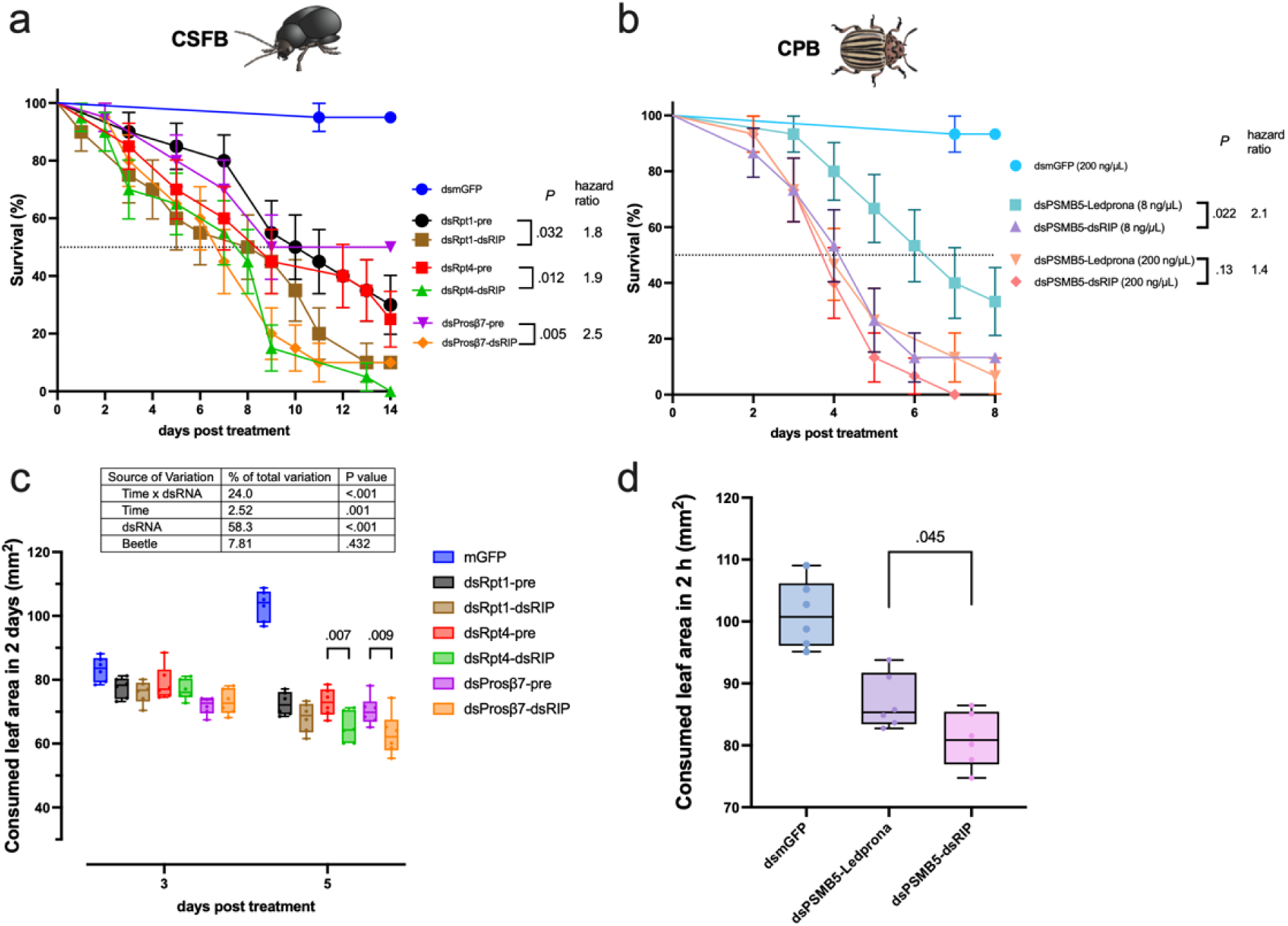
Testing efficacy of double-stranded RNA designed by dsRIP in two leaf beetles. a) *Psylliodes chrysocephala* (cabbage stem flea beetle, CSFB, n = 20 sex-mixed adults per group) was fed with 50 ng of dsRNA used in the previous study or designed using dsRIP against one of the three effective target genes, namely *Pc-rpt4* or *Pc*-*prosB7* and *Pc*-*rpt1* or dsmGFP as control. The daily assessed survival of the adults was plotted (mean survival ± standard error), and the survival curves of two dsRNA designed targeting the same target gene were compared using log-rank test. b) *Leptinotarsa decemlineata* (Colorado potato beetle, CPB, n = 15 L3 larvae) was fed with 600 ng (200 ng/µL * 3 µL) or 24 ng (8 ng/µL * 3 µL) of dsRNA used in commercial product Ledprona or designed using dsRIP against *Ld*-*psmb5* or 600 ng/µL of dsmGFP as control. The daily assessed larval survival was plotted (mean survival ± standard error), and the survival curves of two dsRNA designed targeting the same target gene were compared using log-rank test with Bonferroni correction. c) *P. chrysocephala* (n = 6 sex-mixed adults per group) receiving the 50 ng dsRNA treatments were subjected to feeding area measurements on 3- and 5-days post-treatment, which represented the consumed leaf area in the preceding 2 day period. The data was analyzed through Two-way ANOVA (factors: day and dsRNA) followed by Šídák’s tests between the dsRNAs targeting the same gene. d) *L. decemlineata* (n = 6 L3 larvae per group) receiving the 24 ng dsRNA treatments were subjected to feeding area measurements on the 3 days post-treatment to measure the consumed leaf area in a 2 h period representing the morning activity peak. The data was analyzed through One-way ANOVA followed by Šídák’s test between the dsRNA targeting Ld-*psmb5*.

In *L. decemlineata*, we tested high (600 ng) and low (24 ng) concentrations of the commercialized dsRNA sequence used in Ledprona and dsRNA designed by dsRIP, targeting the same effective target gene *Ld*-*psmb5*, a proteasome subunit (Fig. 4b). Of note, the dsRNA sequence representing Ledprona in this study covered 94% of the actual design used in Ledprona, due to the primer design procedure (see Supporting file 1: *L.decemlineata* dsRNA for details). Although no major difference in insecticidal efficacy was observed at high dsRNA concentration (P = 0.13, hazard ratio: 1.4), dsRIP-designed dsRNA achieved significantly higher performance at the low dsRNA concentration (P = 0.022, hazard ratio: 2.1). We also checked the feeding inhibitory effects of low dsRNA concentrations on 3 days post-treatment (Fig. 4d). dsRNA designed by dsRIP also significantly improved the feeding inhibitory effect (P = 0.045), albeit the difference was marginal: 7% less feeding was observed in dsRIP-designed dsRNA compared to the Ledprona sequence.

The bioassays in two chrysomelids showed that the dsRNA optimization strategy based on siRNA features is applicable across species and also works when dsRNA is delivered orally, which is the primary mode of delivery in RNAi-based pest control. Notably, our fragments performed even somewhat better than the probably extensively optimized commercial sequence. This shows that our design parameters are quite good in predicting effective dsRNA sequences. Given the transferability of our findings, we aimed to make the dsRNA designing strategy available to the community by implementing a new online platform called dsRIP.

### dsRIP web platform

In the first part of this study, we experimentally established a strategy for selecting optimized dsRNA regions with high efficacy based on identified siRNA features. Improving the dsRNA sequence to knock down gene function seems important for both, research and pest control. For pest control, the selection of an effective target gene is essential in the first place and minimizing off-target effects on non-target species is important. To meet the needs for improving any RNAi experiment with a focus on its application to pest control, we developed a web-based platform named dsRIP that integrates tools for a complete *in silico* pipeline for RNAi-based pest control: The target gene finder, the dsRNA efficacy optimizer, the off-target minimizer, and the primer designer (Fig. 5).

**Figure 5.**
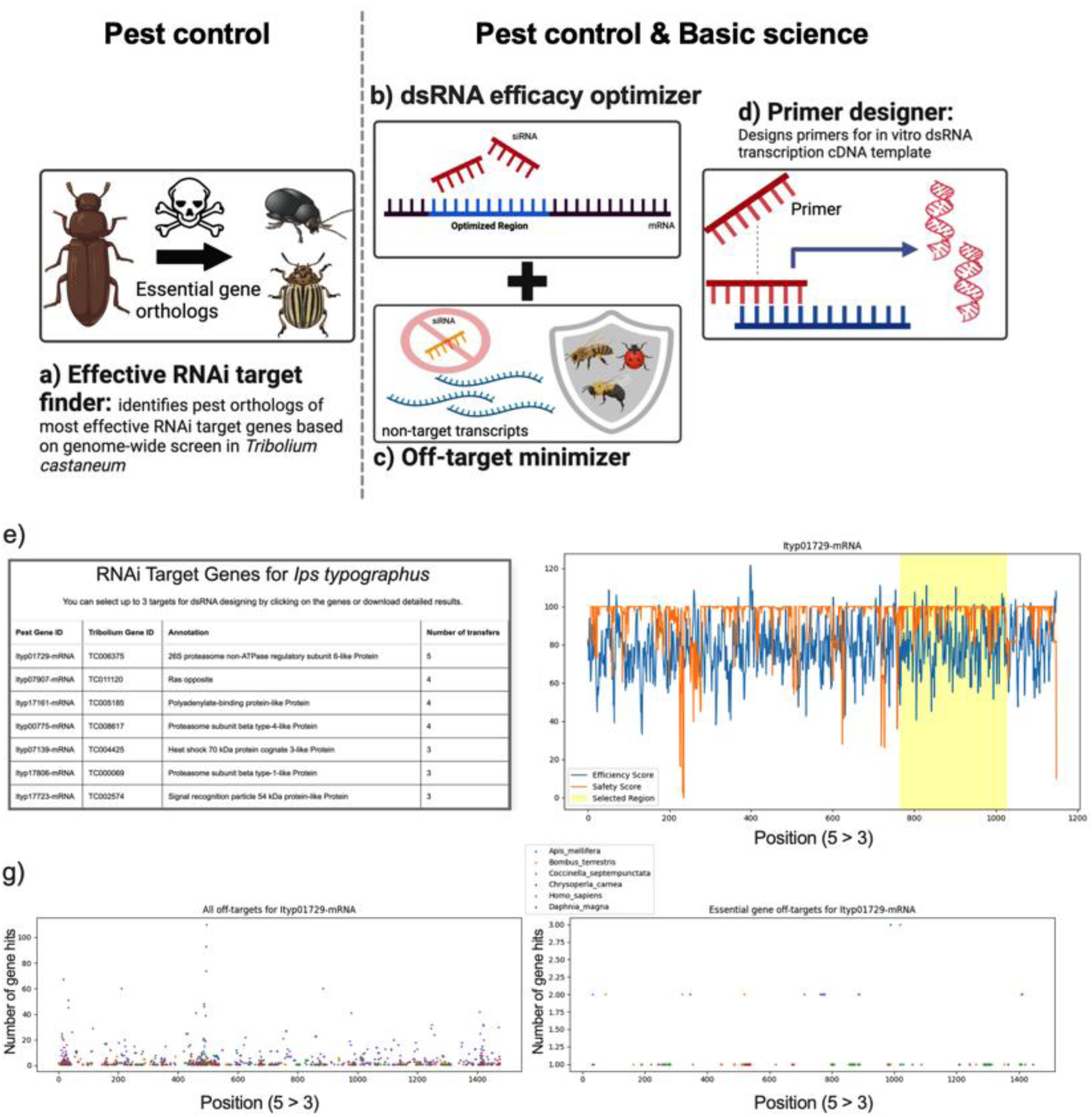
Overview of dsRIP web platform. (a) RNAi target gene finder suggests high-confidence target genes for RNAi-mediated pest control. It finds pest orthologs to the most effective pest control target genes identified through a genome-wide RNAi screen in *T. castaneum*. (b) dsRNA efficacy optimizer uses the findings of the current study to determine the sequence within a target gene with the highest predicted RNAi efficacy. This tool is useful for optimizing RNAi approaches in both basic science and pest control. (c) Off-target minimizer finds sequences within a chosen dsRNA fragment that are highly complementary to sequences in other genes of a given transcriptome. In gene function studies, off-target effects with genes from the studied species need to be avoided. In pest control, the tool increases safety by minimizing harm to selected non-target organisms by calculating a safety score for each region to guide dsRNA designs that are not complementary to transcripts in non-target organisms. (d) dsRIP includes a primer designer tool that can be used to synthesize templates for *in vitro* transcription of the designed dsRNA. (e) Example output with RNAi target genes suggested by dsRIP for *Ips typographus*. (f) An example of a dsRNA design with equal priority to efficacy and safety against the top recommended target gene is in *I. typographus.* The user can choose the balance between efficacy (prioritizing siRNA features) and safety (prioritizing minimization of off-targets) predictions in the final dsRNA design. (g) A detailed safety profile is also provided at the individual non-target organism level. Each dot represents a region that is predicted to have off-target effects on a certain number of genes (Y axis) in the respective species (dot colors) selected by the user. The right graph shows potential off-targets restricted to orthologs of essential genes in the non-target species.

### Effective RNAi target finder

This tool is aimed at users who wish to identify promising RNAi target genes in their pest species. Recently, a genome-wide RNAi screen in the *T. castaneum* was completed, and the most effective target genes were identified [25, 26]. Orthologs of most of these effective targets in *T. castaneum* also achieved high mortality in multiple pest species, namely *Phaedon cochleariae*, *L. decemlineata*, and *P. chrysocephala*. Here, we performed orthology inference using OrthoFinder [45] in many important arthropod and fungi species (around 80 species) and implemented a pipeline for the user to add novel species. The addition of novel species requires the transcriptome or ORF sequences, which can be automatically retrieved from NCBI or uploaded by the user. The transcriptome is converted into protein sequences that are used as an input for orthology inference with *T. castaneum* protein sequences. The curated effective target gene list from Buer et al., together with the orthology information, allows the dsRIP to suggest highly promising RNAi gene targets in the species of interest.

dsRIP provides options for the sorting of candidate target genes based on the number of species in which the orthologous gene was shown to be an effective target. It also allows for the discarding of candidate target genes if they have predicted paralogs, which may functionally compensate for the knocked down of the target gene, thereby reducing insecticidal efficacy. Also, filtering target genes based on their involvement in one of the top pathways identified for efficacious RNAi-mediated pest control is allowed. The proof-of-concept of the RNAi target gene selection tool was recently shown in *P. chrysocephala* in which 11 out of 14 genes suggested by dsRIP methodology achieved significant mortality and 4 very effective target genes were identified [26].

### Off-target minimizer

Different off-target effects have to be considered when using RNAi. When a dsRNA has a sequence stretch >20 bp or so with another gene, then an RNAi treatment can lead to the knock-down of the target and additional genes. This can complicate the interpretation of effects in gene function analyses in basic science. In pest control, off-target refers to the knocking down of essential genes in another species, such that not only the targeted pest but also another species may be affected.

The off-target prediction tool finds potential off-targets in selected transcriptomes by searching for matches between all siRNA that could be generated from the dsRNA and gene sequences in the transcriptomes. As RNAi may tolerate some mismatches, dsRIP allows detection of siRNA with near-perfect matches (1 or 2 mismatches per siRNA) as well. For application in pest control, this could be environmentally important non-target organisms such as honeybees and ladybugs. The program calculates a safety score, which is higher in regions containing less putative off-targets. The non-target organisms to be checked are selected by the user among the supported non-target species. Species not yet supported by dsRIP can be automatically added by providing the name of the species of interest, which prompts the transcriptome to be retrieved from NCBI or by directly uploading a transcriptome file. To focus this analysis on putatively essential genes, the off-target prediction tool is also able to identify whether the potential off-target effect is affecting an ortholog of one of the essential genes identified in *T. castaneum* [25]. This provides a more rational way of minimizing off-targets by considering the gene identity and associated risk rather than considering all potential off-targets equally, which is the approach used in the current off-target prediction methods for RNAi-based pest control. For use in RNAi-based gene function studies, the transcriptome of the same species can be selected to reveal potential genes that are knocked down along with the target and minimize these cases to increase the accuracy of the findings.

The off-target prediction tool is integrated with the efficacy prediction tool such that each siRNA is assigned both safety and efficacy scores, which enables the selection of regions that simultaneously consider both types of predictions (Fig. 5f). To that end, dsRIP has a priority slider that determines coefficients given to efficacy and safety scores. For instance, if the slider is at 50%, efficacy and safety scores will be considered equally, e.g., the average will be calculated. In contrast, if the slider is set to 80% for efficacy, the efficacy score will be considered 4 times as much as the safety score during the dsRNA design.

### Primer design tool

The primer designer tool outputs primers for the amplification of DNA for *in vitro* transcription of the designed dsRNA sequences through the addition of promoter sequences to the primers or for cloning. The primer design tool is based on Primer 3 [46] and attempts to maximize both the amplification of the optimized dsRNA region and effective primer features.

## Discussion

### Increasing efficiency of RNAi-mediated gene function studies

Our results allow us to select effective dsRNA fragments for RNAi experiments and reduce off-target effects. The online tool dsRIP offers an easy way for the community to make use of that knowledge, thereby potentially increasing the sensitivity of many basic science studies, especially in non-model organisms. To the best of our knowledge, this is the only online tool currently available for both purposes. It should be noted that similar tools, primarily for siRNA designing but allowed for dsRNA designing as well, have been published previously, including DEQOR [34] and E-RNAi [47], which are no longer functional online, or si-Fi [48], which never was an online tool. Importantly, the previous tools were based on human siRNA data, while we determined the parameters in an insect pest model. Indeed, some parameters differ between human and insect data, which confirms the validity of our approach.

### Enhanced design of target sequences for pest control

RNAi has great potential in controlling pest populations without having severe detrimental effects on the environment [49, 50]. However, to be a competitive alternative to chemical pesticides, RNAi-based pest control strategies need to be optimized by improving sequence designs and dsRNA formulations [51]. Previous studies highlighted the importance of target gene selection, which remains the primary consideration in pest control. In this study, we empirically tested sequence features to be considered to select the optimal region within the target gene for use in insects. Based on our results and the previous findings, we were able to develop a complete *in silico* pipeline in the form of a web platform for RNAi-based pest control that finds effective target gene orthologs in pests and selects the optimal region within the target gene for high insecticidal efficacy, while minimizing harm to non-target organisms by identifying potential off-targets.

Testing of siRNA-inserted dsRNA suggested that there are major similarities and some differences between the optimal siRNA features for pest control and those identified through therapeutic siRNA optimization efforts in humans. Notably, thermodynamic asymmetry and lack of secondary structure in siRNA were identified as positive predictors of efficacy both here in insects and in human siRNA literature [33]. Such similarities were expected because the biochemical properties of RNA molecules and probably also the RNAi machinery are conserved across species. Our RISC-bound sRNA-seq also suggested that thermodynamic asymmetry impacts the guide strand selection, thereby impacting efficacy. Hence, the mechanism behind the features associated with high efficacy seems to be conserved, at least for thermodynamic asymmetry. Interestingly, higher GC% content between 9^th^ and 14^th^ positions was found to be associated with higher efficacy, contrasting the lower GC% content reported in therapeutic siRNA literature [52, 53]. This could be due to differences in RISC proteins, e.g., the catalytic site of RISC might work more effectively if the local region more strongly hybridizes with the target site in insect pests. These findings support our initial assumption that the previous findings in the therapeutic siRNA field are very useful as a starting point for optimizing RNAi-based pest control but also highlight the fact that these features need to be systematically validated and adjusted for insect species, as discrepancies may also be present.

Designing of dsRNA by considering siRNA-features resulted in improved insecticidal efficacy performance in most cases, which validated our approach. However, it should be noted that some of the insights from siRNA-feature testing experiments could not be well integrated into the dsRNA design. This is because dsRIP treats all potential siRNAs equally, which reduces the significance of specific positional effects, such as the presence of Adenine at the 10th nucleotide, as these features are averaged across all siRNAs considered during dsRNA design. Also, it is still likely that further sequence-features not considered in this study may be associated with RNAi efficacy in insect pests, as our approach did not exhaustively test all possible sequence combinations. Such an exhaustive approach would be necessary to identify efficacy-related features in insects independently of human data, by providing a large dataset for deep learning or similar methodologies. Therefore, further advancements in optimizing dsRNA designs for pest control beyond the strategy established in this study are achievable, and this work may serve as a baseline for future research.

This study also reports a comprehensive data set of RISC-bound siRNA sequencing following the delivery of diverse dsRNA sequences to pest species. It revealed non-random processing of dsRNAs and a preference for the antisense strand as the guide strand for the efficacious dsRNA. Our results also suggest dsRNA processing is probably similar, at least in different coleopteran species (Fig. 3j-k), suggesting that our work in one model pest species such as *T. castaneum* may well generalize to other insect species.

## Conclusion

RNA interference (RNAi) is used widely in basic research and has notable potential as an eco-friendly alternative to chemical pesticides. However, maximum efficacy in research and pest control strategies requires careful sequence design. In this study, we identified key sequence features from insects, such as thermodynamic asymmetry and the lack of secondary structures, that correlate with high insecticidal efficacy. Importantly, we found differences compared to the previous predictors based on human data. Next, we leveraged these findings to improve the insecticidal efficacy of dsRNA in several insect pest species and confirmed that the prediction matched or even exceeded a currently used commercial dsRNA fragment. Furthermore, we developed a comprehensive *in silico* pipeline, available as a web platform named dsRIP (https://dsrip.uni-goettingen.de), which facilitates the identification of effective RNAi targets and the selection of optimal regions for dsRNA designs while minimizing off-target effects in non-target organisms. Overall, the identified features and the developed *in silico* pipeline will enhance the efficacy and safety of RNAi-based pest control strategies. The pipeline will also be valuable to ensure maximum efficacy and accuracy in basic science using RNAi as a tool.

## Methods

### Insect rearing

*Tribolium castaneum* laboratory colony was maintained at 28 °C, 40% relative humidity, under a 16:8 light/dark cycle regime on whole wheat flour. *Psylliodes chrysocephala*, laboratory colony was reared on winter oilseed rape plants (growth stage: BBCH 30–35) at 21° ± 1°C and 65 ± 10% relative humidity under a 16:8 light/dark cycle regime. *Leptinotarsa decemlineata*, was reared on young potato plants at 24° ± 1°C and 65 ± 10% relative humidity under a 16:8 light/dark cycle regime.

### siRNA inserted dsRNA

The essential gene Tc-*gawky* (TC006679, ibeetle-base.uni-goettingen.de) was selected as the target due to being one of the most effective target genes in genome-wide RNAi screen and having a long mRNA (6224 bp), allowing more degrees of freedom in siRNA selection. We designed 31 siRNAs that were 21 nt long and complementary to various regions to the mRNA sequence of Tc-*gawky*. The features of each siRNA were predicted as in dsRNA efficacy optimization tool section using dsRIP. Primers (provided by IDT, Germany) were designed to introduce each siRNA into the GFP sequence modified for no off-targets in *T. castaneum* transcriptome (mGFP; sequence was ordered as gBlock from IDT, Germany) through extension PCR reactions using Q5 High-Fidelity DNA Polymerase (NEB, Germany). The DNA bands with intended insertion were extracted from agarose gels using Gel and PCR Clean-up kit (Macherey-Nagel, Germany) kit. The siRNA insertions were confirmed through similar PCR reactions targeting the siRNA insert. The siRNA inserted mGFP DNA were used for two separate reactions to introduce T7 promoter at the 5’ or 3’ end. Two PCR reactions were conducted for each siRNA inserted dsRNA: one for the sense strand using T7 + forward and reverse primers, and one for the antisense strand using forward and T7 + reverse primers (0.5 μM each). PCRs (50 μL) were carried out with Q5® Hot Start High-Fidelity 2X Master Mix, using an annealing temperature of 60 °C for 28–30 cycles. The expected single band products were confirmed via agarose gel electrophoresis and purified using the Gel and PCR Clean-up kit (Macherey-Nagel, Germany). The sense and antisense RNA strands were synthesized separately in 20 μL reactions using the MEGAscript™ T7 Transcription Kit (Invitrogen, Germany) and purified by Lithium Chloride precipitation. The two RNA strands were combined equimolecularly in nuclease-free water. The RNA strands were annealed by denaturation at 94°C for 5 min, followed by cooling at room temperature for 30 min. The lengths of the annealed dsRNAs were verified on a 1.5% agarose gel alongside a dsRNA ladder (NEB# N0363S, Germany). The siRNA inserted dsRNAs were tested on *T. castaneum* (see Supporting File 1: *T.castaneum* siRNA insert for further details).

### Fully complementary dsRNA

dsRIP was used to select dsRNA regions with different sequence features. For each of the 8 effective target genes from the genome-wide screen in *T. castaneum* [25], we selected a) a region with the highest mean siRNA score within ORF (see dsRNA designer section) and b) a region with lower mean siRNA score within ORF, c) a region with the highest mean accessibility score in whole mRNA, and d) region with the highest mean siRNA score in whole mRNA. The dsRNA regions c or d were not tested if they were greatly overlapping with one of the other regions (>60%). Hence, at least 3 different regions and if possible 4 regions in total were tested for each target gene in *T. castaneum*.

Six different dsRNA targeting two different regions of Pc-*rpt1*, Pc-*rpt4* or Pc-pr*osβ7* were tested in *P. chrysocephala*. One of the tested regions was taken from [26] and the other dsRNA region was designed using dsRIP with the highest mean efficacy siRNA score. In *L. decemlineata*, two dsRNA targeted different regions in Ld-*psmb5*. One of the tested dsRNA was taken from [22] and the other dsRNA was designed using dsRIP with the highest mean siRNA efficacy score.

The primers to obtain cDNA template for in vitro dsRNA transcription were designed using the Primer3 helper function in dsRIP. This function first adds 10 bp buffer region at the 5’ and 3’ end of the dsRNA region and then design specific and effective primers that maximizes the input dsRNA region using Primer3 default settings. Next the T7 promoter sequence (‘GAATTGTAATACGACTCACTATAGG’) was added to the 5′ ends of each primer (provided by IDT, Germany). The DNA template for PCR amplification was prepared from a mix of ten L5 *T. castaneum* larvae, ten 1–25 day-old *P. chrysocephala* adults, or five *L. decemlineata* larvae by RNA extraction using the Quick Tissue Kit (Zymo) and reverse transcription with the LunaScript® RT SuperMix Kit, following the manufacturer’s guidelines. PCRs (50 μL) were carried out with Q5® Hot Start High-Fidelity 2X Master Mix, using an annealing temperature of 58°C–60°C for 28–30 cycles. The expected single band products were confirmed via agarose gel electrophoresis and purified using the Gel and PCR Clean-up kit (Macherey-Nagel, Germany). The dsRNA were synthesized in 20 μL reactions using the MEGAscript™ T7 Transcription Kit (Invitrogen) and purified by Lithium Chloride precipitation. The dsRNA were suspended in nuclease-free water and concentration was measured via Nanodrop (40 μg/OD260). The RNA strands were annealed by denaturation at 94°C for 5 min, followed by cooling at room temperature for 30 min. The annealed dsRNAs (2.5 μg) were verified on a 1.5% agarose gel alongside a dsRNA ladder (NEB# N0363S, Germany).

### Bioassays

Each *T. castaneum* L5 larva was injected with 1 µL of siRNA-inserted dsRNA solution (1000 ng / 1 µL) or 1 µL of fully complementary dsRNA solution (1 ng / 1 µL). After injections, five larvae were placed into each well of 6 well-plates (85.4×127.5 mm, flat bottom). The total number of replications for siRNA-inserted dsRNA bioassays was 20 larvae and it was 25 for fully complementary dsRNA bioassays. In parallel, dsmGFP was injected at the same concentration as control. The larvae were checked daily for mortality with the help of a fine brush over a week.

The bioassays with *P. chrysocephala* were conducted through oral delivery of dsRNA. The dsRNAs were diluted to 50 ng/µL in nuclease-free water and mixed with Triton-X (final concentration 200 ppm, Sigma-Aldrich, Germany). Each Petri dish (60 × 15 mm with vents) was set up as an independent replication, containing one newly emerged adult *P. chrysocephala* and a 30 mm^2^ leaf disk punctured from the first true leaves of oilseed rape plants (growth stage: BBCH ∼35) placed on an agarose gel (1% agarose, 80 mm^3^ volume). Next, 1 μL of the dsRNA solution was evenly spread over the leaf disk’s surface and allowed to dry for 10 minutes before introducing the beetles. After 24 hours, the old agarose gel disks were replaced with untreated 130 mm² leaf disks placed on top of new agarose gel disks and replenishment was performed every third day for the next 14 days. Adult survival was monitored daily. The number of replications for each treatment group was 20 sex-mixed *P. chrysocephala* adults.

The bioassays with *L. decemlineata* were also through oral delivery of dsRNA. Two different concentrations (200 ng/µL and 8 ng/µL) of dsRNAs were prepared in nuclease-free water and mixed with Triton-X (final concentration 200 ppm, Sigma-Aldrich, Germany). Five third instar larvae were placed into each well of 6 well-plates (85.4×127.5 mm, flat bottom) 170 mm^2^ leaf disk punctured from young potato plants placed on an agarose gel (1% agarose, 400 mm^3^ volume). Next, 3 μL of the dsRNA solution was evenly spread over the leaf disk surface and allowed to dry for 10 minutes before introducing the larvae. After 24 hours, the old agarose gel disks were replaced with untreated 400 mm² leaf disks placed on top of new agarose gel disks, and this replacement was repeated every third day over a week. Larval survival was monitored daily. The respective rearing conditions were maintained during the bioassays with the *P. chrysocephala* and *L. decemlineata*.

### RISC-bound small RNA sequencing

Four L5 *T. castaneum* larvae injected with 200 ng of dsRNA (high or low scoring dsRNA targeting one of the eight essential genes or dsmGFP) were sampled 3 days post injection using liquid nitrogen (n = 1-2 per treatment). Also, four *P. chrysocephala* were sampled 3 days post dsmGFP feeding using liquid nitrogen. RNA induced silencing complex (RISC) from the samples were isolated using TraPR Small RNA Isolation Kit according to the manufacturer’s instructions [54]. Briefly, the samples were crushed using RNAse-free micro pestles (Carl Roth GmbH, Germany) in 300 µL lysis buffer (TraPR kit) and homogenized through 20 sec of vortexing. The lysates were clarified trough centrifugation at 10,000 g for 5 min in a centrifuge pre-cooled to +4 °C. The cleared lysates were loaded onto TraPR columns and mixed with the resin by inverting the columns vigorously for 20 sec. The RISC were eluted by centrifugation at 1,000 g for 15 sec and this step was repeated twice with the addition of 250 µL of elution buffer each time into the column to obtain a total of 750 µL RISC containing elution. The RISC-bound small RNA was extracted from the RISC elution through RNA precipitation using a phenol (pH 4.3) / chloroform / isoamylalcohol pre-mix (ROTI®Aqua-P/C/I for RNA extraction, Carl Roth GmbH, Germany). Sodium acetate and carrier substance (TraPR kit) enabled efficient precipitation of small RNA. The pellets were washed thrice using ice-cold 400 µL 80% ethanol and the small RNA were eluted in 10 µL RNA elution buffer.

The libraries from the extracted small RNA were prepared using NEBNext® Multiplex Small RNA Library Prep kit (NEB #E7560S) according to the manufacturer’s instructions. To that end, 6 µL of eluted small RNA (∼500 ng RNA) was used as the RNA input. The 3’ SR Adaptor for Illumina or the 5’ SR Adaptor for Illumina were not diluted as recommended for high small RNA inputs. The final PCR amplification reaction was performed with 11 cycles. The PCR amplified DNA was purified using Monarch PCR&DNA kit (NEB, Germany) using 7:1 binding buffer:sample ratio and DNA eluted in 27.5 µL nuclease free water. The quality of the libraries was checked using an Agilent 2100 Bioanalyzer before and after size selection of 135-150 bp cDNA (corresponding to 16 nt to 30 nt) through 6% PolyAcrylamide gel. The libraries were pooled and sent to BGI for small RNA sequencing on a DNBSEQ-G400 platform. The raw read files were deposited into NCBI Sequence Read Archive (BioProject: PRJNA1157760).

The raw reads were trimmed and quality filtered (>30) using TrimGalore v0.6 (https://github.com/FelixKrueger/TrimGalore) with default parameters. The cleaned reads were mapped onto the respective dsRNA sequences using Bowtie 1 v1.3, which reported all valid mappings in forward and reverse directions without allowing any mismatches (options: “--all”, and “n=0”). The 21 nt-long mapped reads were further analyzed using samtools (v1.2, https://github.com/samtools).

### Leaf consumption

For *P. chrysocephala*, leaf consumption was assessed in two time frames: 1 to 3 days post-treatment and 3 to 5 days post-treatment. Fresh leaves of 130 mm² were provided, and the remaining area was measured two days later (n = 6). For *L. decemlineata*, consumption was measured three days post-treatment by providing fresh leaves of 170 mm², with the remaining area assessed two hours later (n = 6). This measurement period corresponded to the morning activity peak in *L. decemlineata*.

### Statistical analysis

The survival data from bioassays were used to plot Kaplan-Meier survival curves. The siRNA-inserted dsRNA bioassay data was fitted into a Cox regression model, which included the main effects of 9 siRNA sequence features. The fully complementary dsRNA bioassay data obtained in *T. castaneum* was used to calculate the hazard ratio per dsRNA with respect to the dsmGFP control group using the log-rank method. Next, the dsRNA bioassay data was fitted into a multiple regression model, including main effects for mean siRNA score and mean accessibility score, target region and target gene on hazard ratio. The antisense/sense ratios between high and low scoring dsRNA targeting the same gene were analyzed using paired t-test.

The survival curves obtained in bioassays with *P. chrysocephala* and *L. decemlineata* were used to compare previous dsRNA designs with dsRIP-predicted dsRNA designs using one-tailed log-rank test to obtain P value and hazard ratio between the treatments. Bonferroni correction was applied to the P values where applicable. Leaf consumption data for *P. chrysocephala* was analyzed using a two-way ANOVA, with time and dsRNA as factors, followed by Šídák’s multiple comparisons test. For *L. decemlineata*, data was analyzed using a one-way ANOVA, also followed by Šídák’s multiple comparisons test.

### Deployment of dsRIP

The web-based platform dsRIP was developed using Python 3 and Flask library. dsRIP consists of four integrated tools (Tab. 1). dsRIP is hosted by a virtual machine maintained by (GWDG, Göttingen, Germany) that runs Ubuntu v23. dsRIP was deployed using Gunicorn (v22, https://github.com/benoitc/gunicorn) and nginx (https://nginx.org/en/index.html) is used as reverse proxy.

**Table 1.**
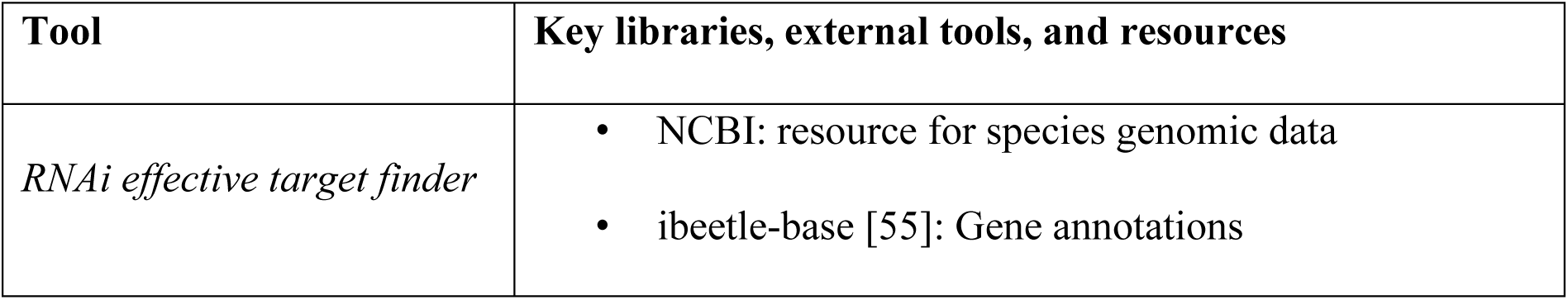
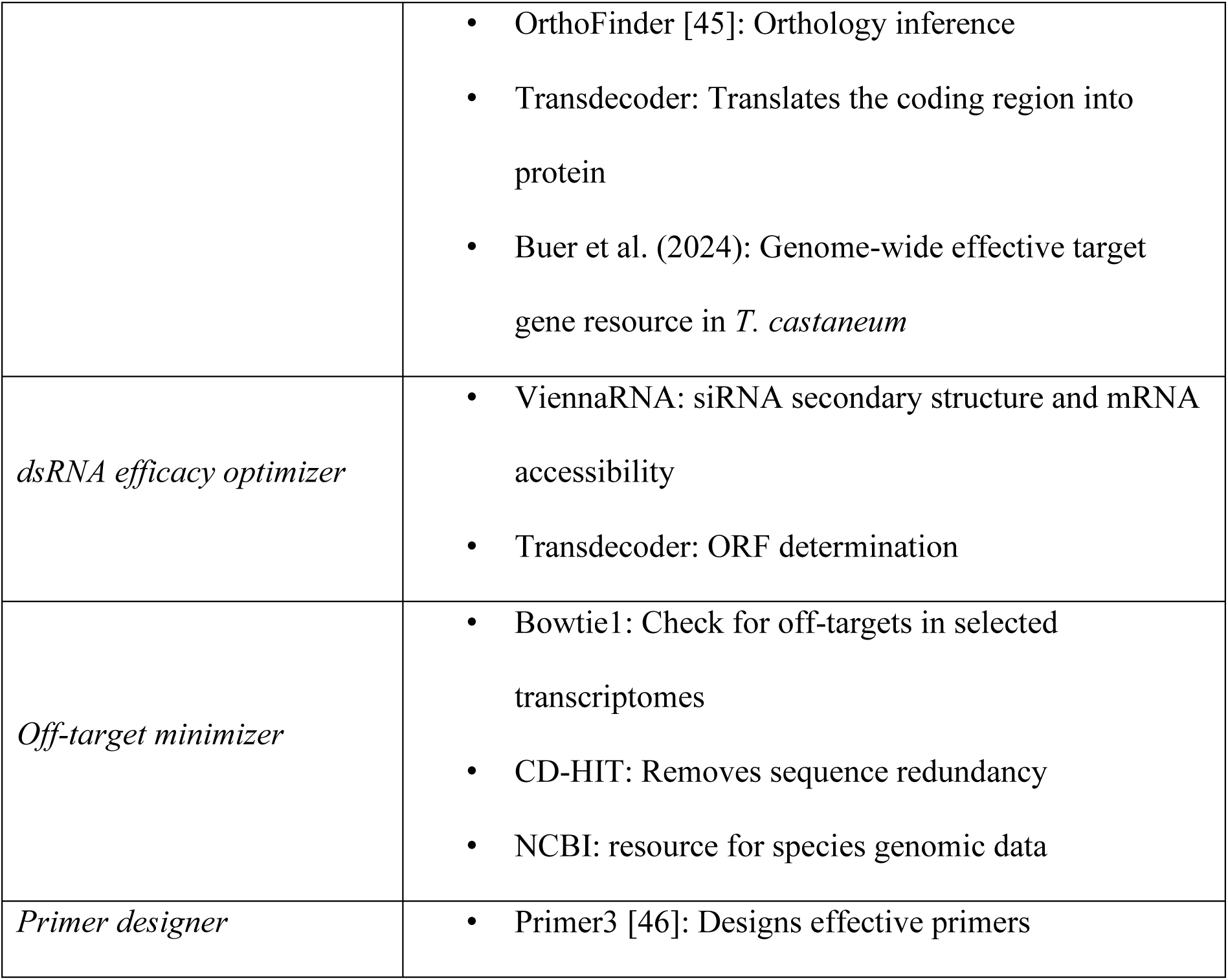
Key libraries and external tools that support the tools in dsRIP.

### RNAi target gene finder

RNAi target gene finder tool filters genes in the selected pest species orthologous to our curated list of most effective target genes [23]. The curated list is based on the genome-wide RNAi screen in *T. castaneum* that identified superior target genes, which were further tested in multiple leaf beetles, including the mustard beetle *Phaedon cochleariae*, Colorado potato beetle *Leptinotarsa decemlineata* and cabbage stem flea beetle *Psylliodes chrysocephala* [25, 26]. OrthoFinder [45] with the default parameters was used to conduct an orthology inference analysis in 83 organisms that showed promise for RNAi-based pest control, other important pests and important non-target animals. For orthology inference, the latest versions of reference transcriptomes or genome extracted coding sequences (if transcriptome was not available) were retrieved from NCBI (https://www.ncbi.nlm.nih.gov/, accessed in May 2024). CD-Hit was used to discard highly similar sequences (>95% identity). TransDecoder (v5.7, https://github.com/TransDecoder/TransDecoder) was used to obtain the longest protein sequences. The same pipeline is used to process user-uploaded transcriptomes.

### dsRNA designer: efficacy and off-target prediction

dsRIP integrates various predictions for each prospective siRNA duplex (21-nt by default) that could be derived from the target gene sequence to calculate efficacy scores and safety scores. The thermodynamic asymmetry between the ends of siRNA strands is calculated to favor the antisense strand in the guide strand selection. Specifically, dsRIP calculates the pairing energy difference between the first four nucleotides of the antisense strand and the last four nucleotides of the sense strand, excluding 2 nucleotide overhangs at the 3’ end of the duplex. We obtained the kcal/mol values for every possible four-base pair long RNA duplex region from a database of RNA duplex stabilities determined by optical melting experiments [56]. dsRIP gives higher scores to siRNA duplexes with weak pairing at the 5’ end of the antisense strand and strong pairing at the 5’ end of the sense strand. We used ViennaRNA v2.7 package to predict the secondary structure and mRNA accessibility of the target region for each prospective siRNA. These quantitative features are percent normalized based on highest and lowest possible values. Next, dsRIP uses the features significantly associated with insecticidal efficacy in this study (shown in Fig. 1c) to calculate an siRNA efficacy score for each prospective siRNA. For the experimental tests in this study, only this siRNA efficacy score was considered during dsRNA designing.

Off-target minimizer tool finds matches between all prospective siRNAs and the transcriptomes of selected non-target organisms using Bowtie1 (v1.3). Bowtie1 is instructed to report all matches regardless of directionality. The number of allowed mismatches for a valid match can be inputted by the user. The found matches for each selected non-target organism are first recorded in “sam” format, which is subsequently parsed to obtain the number of transcript matches for each siRNA. Users can also add their own transcriptomes for off-target identification. The off-target minimizer calculates a safety score for each siRNA by calculating the total number of off-target hits and essential gene off-target hits, with the latter being given a higher coefficient (20:1 by default). The safety score is percent normalized by taking siRNA with the least off-targets as 100% and the one with the most off-targets as 0%. Lastly, the pipeline employs a sliding window to identify the dsRNA region that maximizes the siRNA efficacy score and safety score according to the priority provided by the user.

### Primer designer

Python 3 version of the Primer3 software is implemented to design effective primers for the amplification of the region selected by the dsRNA designer for downstream applications e.g., in vitro transcription. Parameters such as primer annealing temperature and GC content can be provided by the user using the tool’s interface. Also, common promoter sequences can be added to the primers as 5’ overhangs. The output of the primer designer tool is an Excel sheet containing the designed primers per dsRNA.

## ACKNOWLEDGEMENTS

The authors thank GWGD (Gesellschaft für wissenschaftliche Datenverarbeitung mbH Göttingen) personnel for maintaining the virtual machine running the web platform and Stefan Scholten, Shuta Kurokawa, Gerd Vorbrüggen, Claudia Hinners and Jonas Watterott for their help in the project. The project was funded by the Deutscher Akademischer Austauschdienst research grants program. Some of the figures were created with BioRender.com.

## AUTHOR CONTRIBUTION

D.C. and G.B. conceived the ideas and designed the methodology; D.C. performed the experiments in *T. castaneum* and analyzed the data; D.C. and G.G. performed the experiments in two leaf beetles and analyzed the data; D.C. programmed the web platform; G.B. and M.R. provided the workplace and insects. D.C. and M.R. determined the pest and non-target species included in the web platform; D.C. and G.B. drafted and revised the manuscript.

## Data availability

All the primer and template DNA sequences used in this study are included in the supporting file 1. The raw RISC-bound sRNA-seq read files were deposited into NCBI Sequence Read Archive (BioProject: PRJNA1157760). The information for *T. castaneum* genes is publicly available at https://ibeetle-base.uni-goettingen.de. The outputs produced by dsRIP for experimental testing are available on FigShare (https://doi.org/10.6084/m9.figshare.27495843.v1). Any further details can be obtained from the corresponding author.

## Code availability

dsRIP is freely and publicly available as a web-platform at https://dsrip.uni-goettingen.de. Scripts used for RISC-bound sRNA-seq data analysis are available on FigShare (https://doi.org/10.6084/m9.figshare.27490890.v1). Any further details can be obtained from the corresponding author.

## References

1. van Dijk M, Morley T, Rau ML, Saghai Y. A meta-analysis of projected global food demand and population at risk of hunger for the period 2010–2050. Nat Food. 2021;2:494– 501.

2. Gu D, Andreev K, Dupre ME. Major Trends in Population Growth Around the World. China CDC Wkly. 2021;3:604–13.

3. Savary S, Willocquet L, Pethybridge SJ, Esker P, McRoberts N, Nelson A. The global burden of pathogens and pests on major food crops. Nat Ecol Evol. 2019;3:430–9.

4. Farajollahi A, Fonseca DM, Kramer LD, Marm Kilpatrick A. “Bird biting” mosquitoes and human disease: A review of the role of *Culex pipiens* complex mosquitoes in epidemiology. Infection, Genetics and Evolution. 2011;11:1577–85.

5. Sparks TC, Nauen R. IRAC: Mode of action classification and insecticide resistance management. Pestic Biochem Physiol. 2015;121:122–8.

6. Chen YH, Cohen ZP, Bueno EM, Christensen BM, Schoville SD. Rapid evolution of insecticide resistance in the Colorado potato beetle, Leptinotarsa decemlineata. Curr Opin Insect Sci. 2023;55:101000.

7. Krattinger SG, Keller B. Molecular genetics and evolution of disease resistance in cereals. New Phytologist. 2016;212:320–32.

8. Liu N. Insecticide resistance in mosquitoes: impact, mechanisms, and research directions. Annu Rev Entomol. 2015;60:537–59.

9. Serrão JE, Plata-Rueda A, Martínez LC, Zanuncio JC. Side-effects of pesticides on non-target insects in agriculture: a mini-review. Sci Nat. 2022;109:17.

10. Devi PI, Manjula M, Bhavani RV. Agrochemicals, Environment, and Human Health. Annual Review of Environment and Resources. 2022;47 Volume 47, 2022:399–421.

11. Nicholson CC, Knapp J, Kiljanek T, Albrecht M, Chauzat M-P, Costa C, et al. Pesticide use negatively affects bumble bees across European landscapes. Nature. 2024;628:355–8.

12. Baum JA, Bogaert T, Clinton W, Heck GR, Feldmann P, Ilagan O, et al. Control of coleopteran insect pests through RNA interference. Nat Biotechnol. 2007;25:1322–6.

13. Huvenne H, Smagghe G. Mechanisms of dsRNA uptake in insects and potential of RNAi for pest control: A review. Journal of Insect Physiology. 2010;56:227–35.

14. Meister G, Tuschl T. Mechanisms of gene silencing by double-stranded RNA. Nature. 2004;431:343–9.

15. Fire A, Xu S, Montgomery MK, Kostas SA, Driver SE, Mello CC. Potent and specific genetic interference by double-stranded RNA in Caenorhabditis elegans. Nature. 1998;391:806–11.

16. Koo J, Palli SR. Recent advances in understanding of the mechanisms of RNA interference in insects. Insect Molecular Biology. n/a n/a.

17. Koo J, Palli SR. StaufenC facilitates utilization of the ERAD pathway to transport dsRNA through the endoplasmic reticulum to the cytosol. Proceedings of the National Academy of Sciences. 2024;121:e2322927121.

18. Zhu KY, Palli SR. Mechanisms, Applications, and Challenges of Insect RNA Interference. Annual Review of Entomology. 2020;65 Volume 65, 2020:293–311.

19. Preall JB, Sontheimer EJ. RNAi: RISC Gets Loaded. Cell. 2005;123:543–5.

20. Orban Ti, Izaurralde E. Decay of mRNAs targeted by RISC requires XRN1, the Ski complex, and the exosome. RNA. 2005;11:459–69.

21. Reinders JD, Moar WJ, Head GP, Hassan S, Meinke LJ. Effects of SmartStax® and SmartStax® PRO maize on western corn rootworm (Diabrotica virgifera virgifera LeConte) larval feeding injury and adult life history parameters. PLoS One. 2023;18:e0288372.

22. Rodrigues TB, Mishra SK, Sridharan K, Barnes ER, Alyokhin A, Tuttle R, et al. First Sprayable Double-Stranded RNA-Based Biopesticide Product Targets Proteasome Subunit Beta Type-5 in Colorado Potato Beetle (Leptinotarsa decemlineata). Frontiers in Plant Science. 2021;12.

23. Cedden D, Bucher G. The quest for the best target genes for RNAi-mediated pest control. Insect Molecular Biology. n/a n/a.

24. Mehlhorn S, Hunnekuhl VS, Geibel S, Nauen R, Bucher G. Establishing RNAi for basic research and pest control and identification of the most efficient target genes for pest control: a brief guide. Front Zool. 2021;18:60.

25. Buer B, Dönitz J, Milner M, Mehlhorn S, Hinners C, Siemanowski-Hrach J, et al. Superior target genes and pathways for RNAi mediated pest control revealed by genome wide analysis in the red flour beetle Tribolium castaneum. 2024;:2024.01.24.577003.

26. Cedden D, Güney G, Debaisieux X, Scholten S, Rostás M, Bucher G. Effective target genes for RNA interference-based management of the cabbage stem flea beetle. 2024;:2024.04.30.591975.

27. Hurowitz EH, Brown PO. Genome-wide analysis of mRNA lengths in Saccharomyces cerevisiae. Genome Biol. 2003;5:R2.

28. Wang K, Peng Y, Fu W, Shen Z, Han Z. Key factors determining variations in RNA interference efficacy mediated by different double-stranded RNA lengths in Tribolium castaneum. Insect Molecular Biology. 2019;28:235–45.

29. Bolognesi R, Ramaseshadri P, Anderson J, Bachman P, Clinton W, Flannagan R, et al. Characterizing the Mechanism of Action of Double-Stranded RNA Activity against Western Corn Rootworm (Diabrotica virgifera virgifera LeConte). PLOS ONE. 2012;7:e47534.

30. He W, Xu W, Xu L, Fu K, Guo W, Bock R, et al. Length-dependent accumulation of double-stranded RNAs in plastids affects RNA interference efficiency in the Colorado potato beetle. Journal of Experimental Botany. 2020;71:2670–7.

31. Kurreck J. siRNA Efficiency: Structure or Sequence—That Is the Question. J Biomed Biotechnol. 2006;2006:83757.

32. Reynolds A, Leake D, Boese Q, Scaringe S, Marshall WS, Khvorova A. Rational siRNA design for RNA interference. Nat Biotechnol. 2004;22:326–30.

33. Naito Y, Yoshimura J, Morishita S, Ui-Tei K. siDirect 2.0: updated software for designing functional siRNA with reduced seed-dependent off-target effect. BMC Bioinformatics. 2009;10:392.

34. Henschel A, Buchholz F, Habermann B. DEQOR: a web-based tool for the design and quality control of siRNAs. Nucleic Acids Research. 2004;32 suppl_2:W113–20.

35. Noland CL, Ma E, Doudna JA. siRNA Repositioning for Guide Strand Selection by Human Dicer Complexes. Molecular Cell. 2011;43:110–21.

36. Schwarz DS. Asymmetry in the assembly of the RNAi enzyme complex. Cell. 2003;115.

37. Tants J-N, Fesser S, Kern T, Stehle R, Geerlof A, Wunderlich C, et al. Molecular basis for asymmetry sensing of siRNAs by the Drosophila Loqs-PD/Dcr-2 complex in RNA interference. Nucleic Acids Res. 2017;45:12536–50.

38. Tomari Y, Matranga C, Haley B, Martinez N, Zamore PD. A protein sensor for siRNA asymmetry. Science. 2004;306.

39. Cedden D, Güney G, Scholten S, Rostás M. Lethal and sublethal effects of orally delivered double-stranded RNA on the cabbage stem flea beetle, Psylliodes chrysocephala. Pest Management Science. 2024;80:2282–93.

40. He W, Xu W, Fu K, Guo W, Kim DS, Zhang J. Positional effects of double-stranded RNAs targeting β*-Actin* gene affect RNA interference efficiency in Colorado potato beetle. Pesticide Biochemistry and Physiology. 2022;184:105121.

41. Tomoyasu Y, Miller SC, Tomita S, Schoppmeier M, Grossmann D, Bucher G. Exploring systemic RNA interference in insects: a genome-wide survey for RNAi genes in Tribolium. Genome Biology. 2008;9:R10.

42. Cooper AM, Silver K, Zhang J, Park Y, Zhu KY. Molecular mechanisms influencing efficiency of RNA interference in insects. Pest Management Science. 2019;75:18–28.

43. J P, A F. Distinct populations of primary and secondary effectors during RNAi in C. elegans. Science (New York, NY). 2007;315.

44. Santos D, Mingels L, Vogel E, Wang L, Christiaens O, Cappelle K, et al. Generation of Virus- and dsRNA-Derived siRNAs with Species-Dependent Length in Insects. Viruses. 2019;11.

45. Emms DM, Kelly S. OrthoFinder: phylogenetic orthology inference for comparative genomics. Genome Biology. 2019;20:238.

46. Untergasser A, Cutcutache I, Koressaar T, Ye J, Faircloth BC, Remm M, et al. Primer3— new capabilities and interfaces. Nucleic Acids Research. 2012;40:e115.

47. Horn T, Boutros M. E-RNAi: a web application for the multi-species design of RNAi reagents—2010 update. Nucleic Acids Research. 2010;38 suppl_2:W332–9.

48. Lück S, Kreszies T, Strickert M, Schweizer P, Kuhlmann M, Douchkov D. Frontiers | siRNA-Finder (si-Fi) Software for RNAi-Target Design and Off-Target Prediction. 10.3389/fpls.2019.01023.

49. Castellanos NL, Smagghe G, Taning CNT, Oliveira EE, Christiaens O. Risk assessment of RNAi-based pesticides to non-target organisms: Evaluating the effects of sequence similarity in the parasitoid wasp *Telenomus podisi*. Science of The Total Environment. 2022;832:154746.

50. Chen Y, De Schutter K. Biosafety aspects of RNAi-based pests control. Pest Management Science. 2024;80:3697–706.

51. Quilez-Molina AI, Niño Sanchez J, Merino D. The role of polymers in enabling RNAi-based technology for sustainable pest management. Nat Commun. 2024;15:9158.

52. Hu B, Zhong L, Weng Y, Peng L, Huang Y, Zhao Y, et al. Therapeutic siRNA: state of the art. Sig Transduct Target Ther. 2020;5:1–25.

53. Friedrich M, Aigner A. Therapeutic siRNA: State-of-the-Art and Future Perspectives. BioDrugs. 2022;36:549–71.

54. Grentzinger T, Oberlin S, Schott G, Handler D, Svozil J, Barragan-Borrero V, et al. A universal method for the rapid isolation of all known classes of functional silencing small RNAs. Nucleic Acids Res. 2020;48:e79.

55. Dönitz J, Schmitt-Engel C, Grossmann D, Gerischer L, Tech M, Schoppmeier M, et al. iBeetle-Base: a database for RNAi phenotypes in the red flour beetle Tribolium castaneum. Nucleic Acids Research. 2015;43:D720–5.

56. Zuber J, Schroeder SJ, Sun H, Turner DH, Mathews DH. Nearest neighbor rules for RNA helix folding thermodynamics: improved end effects. Nucleic Acids Research. 2022;50:5251–62.

