## Supporting information 1 for "Optimizing dsRNA sequences for RNAi in pest control and research with the dsRIP Web-Platform"

**Table S1. Scores and hazard ratios of dsRNA tested in *Tribolium castaneum***

| Target gene | siRNA score | Accessibility score | Region | Hazard ratio |
|---|---|---|---|---|
| ***Tc-gawky*** | 74.3 | 73.8 | ORF | 36.8 |
| ***Tc-gawky*** | 66.4 | 67.4 | ORF | 5.75 |
| ***Tc-gawky*** | 71.4 | 85.4 | ORF | 23.2 |
| ***Tc-gawky*** | 78.4 | 78.5 | UTR | 9.78 |
| ***Tc-klp61F*** | 76.9 | 78 | ORF | 12.1 |
| ***Tc-klp61F*** | 69.7 | 70 | ORF | 4.88 |
| ***Tc-klp61F*** | 73.3 | 86.4 | ORF | 9.9 |
| ***Tc-klp61F*** | 77.8 | 78.7 | UTR | 10.7 |
| ***Tc-nito*** | 77 | 71.6 | ORF | 29.4 |
| ***Tc-nito*** | 65.3 | 73.1 | ORF | 3 |
| ***Tc-nito*** | 74.8 | 78.9 | UTR | 6.34 |
| ***Tc-nito*** | 78.2 | 74.6 | UTR | 12.4 |
| ***Tc-hr3*** | 75.1 | 75.1 | ORF | 11.1 |
| ***Tc-hr3*** | 72 | 72.1 | ORF | 1.55 |
| ***Tc-hr3*** | 76.3 | 80.4 | UTR | 5.17 |

| | | | | |
|---|---|---|---|---|
| ***Tc-hr3*** | 77.8 | 76.6 | UTR | 11.8 |
| ***Tc-eIF3a*** | 75.3 | 66 | ORF | 6.5 |
| ***Tc-eIF3a*** | 65.1 | 58.1 | ORF | 4.14 |
| ***Tc-eIF3a*** | 73.4 | 80.5 | ORF | 5.39 |
| ***Tc-rpt1*** | 74.3 | 74.7 | ORF | 5.29 |
| ***Tc-rpt1*** | 67.6 | 57.6 | ORF | 4.45 |
| ***Tc-rpt1*** | 72.8 | 78.3 | ORF | 5.75 |
| ***Tc-rpn7*** | 76.1 | 79.2 | ORF | 10.4 |
| ***Tc-rpn7*** | 71 | 76.6 | ORF | 8.02 |
| ***Tc-rpn7*** | 76.2 | 76.4 | UTR | 6.14 |
| ***Tc-cyp4g15*** | 73.57 | 73.96 | ORF | 12.7 |
| ***Tc-cyp4g15*** | 69.47 | 69.5 | ORF | 11.9 |
| ***Tc-cyp4g15*** | 71.6 | 78.1 | ORF | 12.6 |

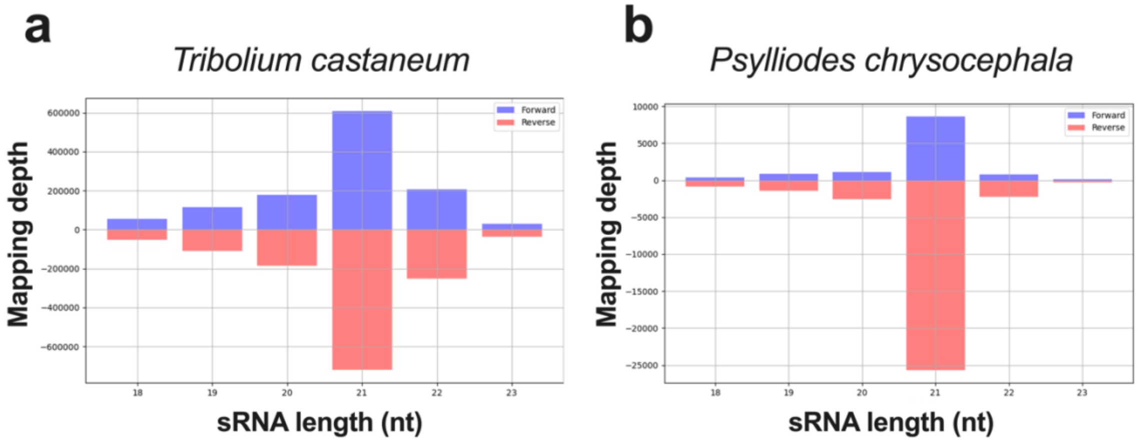


**Figure S1. Length distribution in RISC-bound sRNA mapping to the respective dsRNA sequence.** a) All RISC-bound sRNA-seq experiments in *T. castaneum* larvae were combined to plot the overall length distribution (n = 16). b) The RISC-bound sRNA length distribution in *P. chrysocephala* adults fed on dsmGFP were sampled for RISC-bound sRNA-seq.

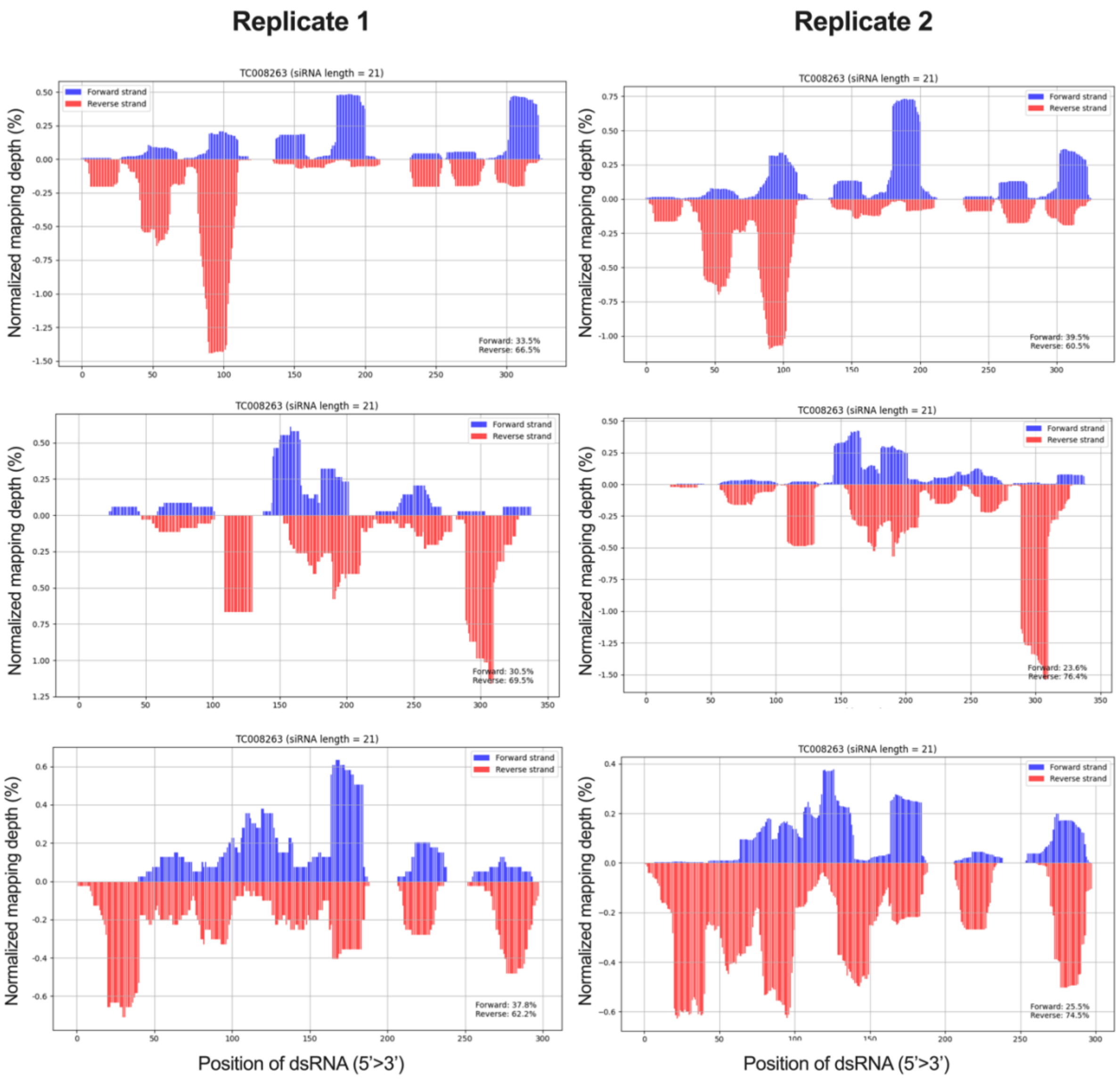


**Figure S2. Reproducibility of RISC-bound sRNA-seq experiments.** RISC-bound sRNA-seq was performed after the delivery of dsRNA targeting the essential gene Tc-*klp61F* (TC008263) to *T. castaneum* L5 larvae. The mapping depths obtain in two biological replications for each dsRNA are plotted separately (left and right plots).
